# Unveiling the hidden electroencephalographical rhythms during development: aperiodic and periodic activity in healthy subjects

**DOI:** 10.1101/2024.05.29.596404

**Authors:** Brenda Y. Angulo-Ruiz, Elena I. Rodríguez-Martínez, Vanesa Muñoz, Carlos M. Gómez

## Abstract

**Objective:** The study analyzes power spectral density (PSD) components, aperiodic (AP) and periodic (P) activity, in resting-state EEG of 240 healthy subjects from 6 to 29 years old, divided into 4 groups.

**Methods:** We calculate AP and P components using the (*Fitting Oscillations and One-Over-f (FOOOF))* plugging in EEGLAB. All PSD components were calculated from 1-45Hz. Topography analysis, Spearman correlations, and regression analysis with age were computed for all the components.

**Results:** AP and P activity show different topography across frequencies and age groups. Age-related decreases in AP exponent and offset parameters lead to reduced power, while P power decreases (1-6Hz) and increases (10-15Hz) with age.

**Conclusions:** We support the distinction between the AP and P components of the PSD and its possible functional changes with age. AP power is dominant in the configuration of the canonical EEG rhythms topography, although P contribution to topography is embedded in the canonical EEG topography. Some EEG canonical characteristics are similar to those of P component, as topographies of EEG rhythms (embedded) and increases in oscillatory frequency with age.

**Significance:** We support that spectral power parameterization improves the interpretation and neurophysiological and functional accuracy of brain processes.

**Highlights:**

- Differential topography for aperiodic and periodic EEG components.
- Power decrease in aperiodic component (exponent and offset) with age.
- Decrease (lower frequencies) and increase (higher frequencies) of periodic component.

## 1. Introduction

Extensive research on neural oscillations has shown how they can be related to a wide variety of cognitive and behavioral states (Engel et al., 2001; Buzsáki and Draguhn, 2004; Donoghue et al., 2022). The transfer of interregional brain information would be one of the most important functions of neural oscillations (Fries, 2005; Voytek et al., 2015a). Recent research suggests a major limitation in using a canonical analysis of predefined frequency bands, known as delta, theta, alpha, beta, and gamma, given the presence of 1/f ^x^ noise embedded with genuine brain rhythms (Donoghue et al., 2020a; He, 2014; Ostlund et al., 2022). An analysis to overcome the latter limitation in the analysis of electrophysiological neural activity, which separates the power spectral density (PSD) into aperiodic (AP) (1/f-like) and periodic (P) activity components has been proposed. AP and P components have independent physiological information, potentially leading to a loss of information in the physiological interpretations of cognitive and behavioral states when analyzing spectral power without differentiating AP and P EEG components (Donoghue et al., 2020a; McSweeney et al., 2023; Podvalny et al., 2015).

The AP component (1/f^x^) is characterized by offset and exponent parameters, showing a decrease in spectral power with increasing frequencies (Donoghue et al., 2020a). The offset parameter, which reflects the change in broadband power across frequencies (Hill et al., 2022), has been associated with increases in neuronal spiking of cortical neurons (Manning et al., 2009; Miller et al., 2014; Voytek and Knight, 2015). The exponent parameter, which reflects the intensity of the power spectrum decay (Donoghue et al., 2020a), has been linked to the integration/balance of excitatory (E) and inhibitory (I) underlying synaptic currents (Buzsáki et al., 2012; Gao et al., 2017). Thus, a lower exponent (flatter PSD) indicated larger excitatory currents compared to inhibitory ones (E>I), while a higher exponent suggests the opposite (Donoghue et al., 2020a). The exponent indicates the maturation of the E/I balance (Cellier et al., 2021).

The balance of E/I currents during development would reflect changes in myelination, cortical volume, neuroreceptors, neurotransmitters, and cortical thickness, all of which are crucial for typical cortical development and function (Donoghue et al., 2020a). Indeed, alterations in this E/I balance have been observed in neurodevelopmental disorders (Levin et al., 2020; Mamiya et al., 2021; Ostlund et al., 2021; Robertson et al., 2019) and psychiatric disorders (Molina et al., 2020; Wilkinson and Nelson, 2021). Furthermore, growing evidence for the presence of AP in PSD suggests changes in AP PSD during development and typical aging (He et al., 2019; McSweeney et al., 2021, 2023; Schaworonkow and Voytek, 2021; Voytek et al., 2015b). Considering the decrease of exponent and offset in the AP component as indicative of optimal brain maturation or development (Cellier et al., 2021; Donoghue et al., 2020a; Hill et al., 2022; McSweeney et al., 2021, 2023; Schaworonkow and Voytek, 2021).

On the other hand, AP adjusted or parameterized P activity is defined by parameters such as center (or peak) frequency, power, and bandwidth; and suggests a possible difference between a traditional spectral power approach and a genuine oscillatory power approach (Donoghue et al., 2020a; McSweeney et al., 2023). In contrast to the latter, classic spectral analysis would combine analysis of P and AP PSD components, biasing physiological interpretations of cognition and behavior (Donoghue et al., 2020a; Ostlund et al., 2022). The characterization of P activity is, therefore, delimited by narrowband power peaks above the AP component, and would describe the genuine rhythmic components of the power spectrum (Donoghue et al., 2020a). Investigations of the different peaks of P show age-related changes during development (Cellier et al., 2021; Hill et al., 2022; Ostlund et al., 2022). These changes include the age-related increase in peak frequency of 4-12Hz oscillations (Cellier et al., 2021), coincident with canonical bands such as theta and alpha, or alpha alone (Cellier et al., 2021; McSweeney et al., 2023), as well as in the beta peak oscillation (He et al., 2019). All of these changes are significant for cognitive development (Cellier et al., 2021).

As can be seen, parameterization points towards the existence of different components in the EEG, which may contribute differentially to cognition across development. Therefore, this study aims to characterize the differences between PSD components and their developmental changes during maturation, by separating the AP and P components of spectral power and analyzing their differences in terms of topography, age maturation, and relationship to behavior. Differences in topography would suggest different neural sources for AP and P components but also would indicate frequency changes with age in defined cortical areas (Rodríguez-Martinez et al., 2017). Employing age as an explanatory variable of the dynamic age-related changes in the AP and P components would complement previous studies. While previous studies have explored the age-related aspects of these components (Donoghue et al., 2020a; Cellier et al., 2021; Schaworonkow and Voytek, 2021; Hill et al., 2022; McSweeney et al., 2021, 2023), they have not explored the effect of age across the whole scalp in AP and P, in a sample covering a so broad age range as in the present report with respect EEG rhythms topographies (6-29 years). Although the relationship with behavioral variables of AP and P components has been explored (Dave et al., 2018; Donoghue et al., 2020a; Gao et al., 2017; Mamiya et al., 2021; Ostlund et al., 2021; Tran et al., 2020), in the present report we would test the possible influence of AP and P components in working memory development.

## 2. Methods

### 2.1. Sample

A total of 240 healthy human participants aged 6-29 years (M=14.38 years, SD=6.03, 117 females) were included. Participants did not report any neurological, psychological illness, or cognitive impairment. The recordings of some subjects included in the analyzed data have been previously employed (Rodríguez-Martínez et al., 2020; Angulo-Ruiz et al., 2021) with different purposes using the classical (canonical) PSD analysis without separating P and AP EEG components. Subjects were grouped into four groups of different age ranges: 6-9 years (63 subjects), 10-13 years (59 subjects), 14-17 years (48 subjects) and 18-29 years (70 subjects) for the analysis.

The sample of children had a normal academic record and the young adults were university students. The study was conducted following the guidelines of the Declaration of Helsinki, with the written informed consent for the adults, and with parental consent for the children sample. The experimental protocol was approved by the biomedical research ethics committee of the Autonomous Community of Andalucía.

### 2.2. Psychological Measures

Different psychological measures of the subjects were independently recorded. These were the mean reaction time (RT) of the visual oddball task (for more detail Rojas-Benjumea et al., 2015); Working Memory Test Battery for Children (*WMTBC;* Pickering and Gathercole, 2001) with its subtests (Phonological Loop (PL), Visuospatial Sketchpad (VSS) and Central Executive (CE)) (for more detail Gómez et al., 2018). It should be noted that there are some missing values (Table 1) due to some subjects did not complete some of the tasks or did not return another day to finish the assessment. U Mann-Whitney test showed no significant difference between the psychological measures for gender after the false discovery rate (FDR) correction (Benjamini and Hochberg, 1995).

**Table 1.** Frequencies and Descriptive analysis of behavioral measures. N=number of subjects, Missing=missing values, M=Media, SD=Standard Deviation

| Test | N | Missing | M | SD |
| --- | --- | --- | --- | --- |
| RT | 237 | 3 | 446.73 | 63.86 |
| LP | 238 | 2 | 121.64 | 19.504 |
| VSS | 238 | 2 | 52.91 | 9.53 |
| CE | 238 | 2 | 71.85 | 15.99 |

### 2.3. EEG recording

The spontaneous recording of EEG brain electrical activity with open eyes had a duration of 3 minutes. Subjects were asked to look at a cross on the screen and blink as little as possible while remaining calm and maintaining a comfortable position. The recording was obtained from 32 electrodes mounted on an electrode cap (ELECTROCAP) selected from the international 10-20 system (Fp1, Fpz, Fp2, F7, F3, Fz, F4, F8, FC5, FC1, FC2, FC6, M1, T7, C3, Cz, C4, T8, M2, CP5, CP1, CP2, CP6, P7, P3, Pz, P4, P8, POz, O1, Oz, O2). Eye movements were recorded by two electrodes placed at the outer edge of each eye for horizontal movements, and two electrodes were placed above and below the left eye for vertical movements. For signal analysis, the ocular and mastoid electrodes were removed and an average reference was used, resulting in a total of 30 analyzed electrodes. Impedance was kept below 10 kΩ. Data were recorded in direct current at 512Hz, with an amplification gain of 20,000 using an analog-digital acquisition and analysis system (ANT Amplifiers, The Netherlands). Data were not filtered during recording.

### 2.4. Data Analysis

#### 2.4.1. EEG pre-processing

Unfiltered EEG data were analyzed with the software package EEGLAB (Delorme and Makeig, 2004) and Matlab R2021b. Signal pre-processing consisted of applying (i) a 47-53Hz notch filter (EEGLAB function: *eegfiltnew*), (ii) an average reference, and (iii) an artifact subspace reconstruction (*ASR*) algorithm (EEGLAB function*: clean raw data*) to correct portions of the data with a standard deviation greater than 20 times that of the calibration data (Mullen et al., 2015). Artifacts from the eyes, blinks, muscles, and other movements were removed using independent component analysis (*ICA*; function: *pop_runica*) with the natural gradient function (Bell and Sejnowski, 1995; Amari et al., 1995), and the ICLabel-extension classification (Pion-Tonachini et al., 2019) in EEGLAB. The EEG signal was reconstructed and all epochs (2000ms duration) in which the EEG exceeded ±120 μV in any electrode, were rejected (function *eegthresh*).

Subjects had a range of selected epochs between 44 and 90 for analysis (M=89.15, SD=3.854), and a range of 20 to 23 accepted components (M=21.63, SD=0.601). Kruskal-Wallis test revealed significant differences in the epochs (H(3, N=240) =54.37, p<.001, ε^2^=0.23) and components (H(3, N=240) =26.89, p<.001, ε^2^=0.11) within the four age groups. The mean rank of the epochs was 84.32 for group 1, 119.34 for group 2, 133.78 for group 3, and 144.68 for group 4. For the components the mean rank was 148.60 for group 1, 129.53 for group 2, 99.25 for group 3 and 102.18 for group 4. Post hoc comparisons using the Dunn method with Bonferroni correction indicated that the mean rank of trials of group 1 was significantly lower than group 2 (p<.001), group 3 (p<.001) and group 4 (p<.001), as well as group 2 compared to group 4 (p=.024). For the components, the mean rank of group 1 was significantly higher than the mean rank of group 3 (p<.001) and group 4 (p<.001). U Mann-Whitney test showed no significant differences in epochs and components with gender.

#### 2.4.2. Parameterization of Fitting Oscillations and One-Over-f (FOOOF)

The full model fit (PSD adjusted), empirical PSD as well as AP and P activity using the *specparam* plugging from FOOOF (https://fooof-tools.github.io/fooof/reference.html) were computed. This plugging uses a MATLAB-adapted version of Python’s *Fitting Oscillations and One-Over-f* (FOOOF) function (version 3.8) (Donoghue et al., 2020a). *Specparam* defines the power spectrum as a combination of AP (in log-log space) and P (oscillations above AP activity) components.

The PSD AP component was calculated using the offset and exponent parameters generated by the *FOOOF* algorithm and was fitted to the entire range of the spectrum. The P component is generated by oscillatory peaks above the AP component, modeled by a Gaussian and whose three parameters (peak power, center frequency, and bandwidth) characterize the oscillation (for more details see Donoghue et al., 2020a). Since the FOOOF algorithm does not return the spectrum-adjusted P component, we computed this component by subtracting the AP component (ap_fit in the *specparam* function) from the PSD adjusted (Figure 8A):

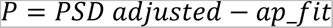

The *FOOOF* function calculates the power spectra for all electrodes and subjects using the Welch method of *spectopo* function in EEGLAB. Recommended algorithm settings were set as: peak_width_limits=[1,8], min_peak_height=0.05, peak_threshold=0.5, max_n_peaks=6, aperiodic_mode= ‘fixed’ (Donoghue et al., 2020a; McSweeney et al., 2023). PSD values were computed from 1 to 45Hz. The variance explained (R^2^) and mean absolute error (MAE) metrics assessed the goodness-of-fit measures between the model and the original power spectrum. Based on the values reported by Ostlund et al. (2022) (MAE underfit>0.1, MAE overfit<0.020), Supplementary Figure 1 shows a good fit of the algorithm in each of the segmented groups in our data. U Mann-Whitney test showed that there were no significant differences in gender for the exponent and offset parameters.

**Figure 1.**
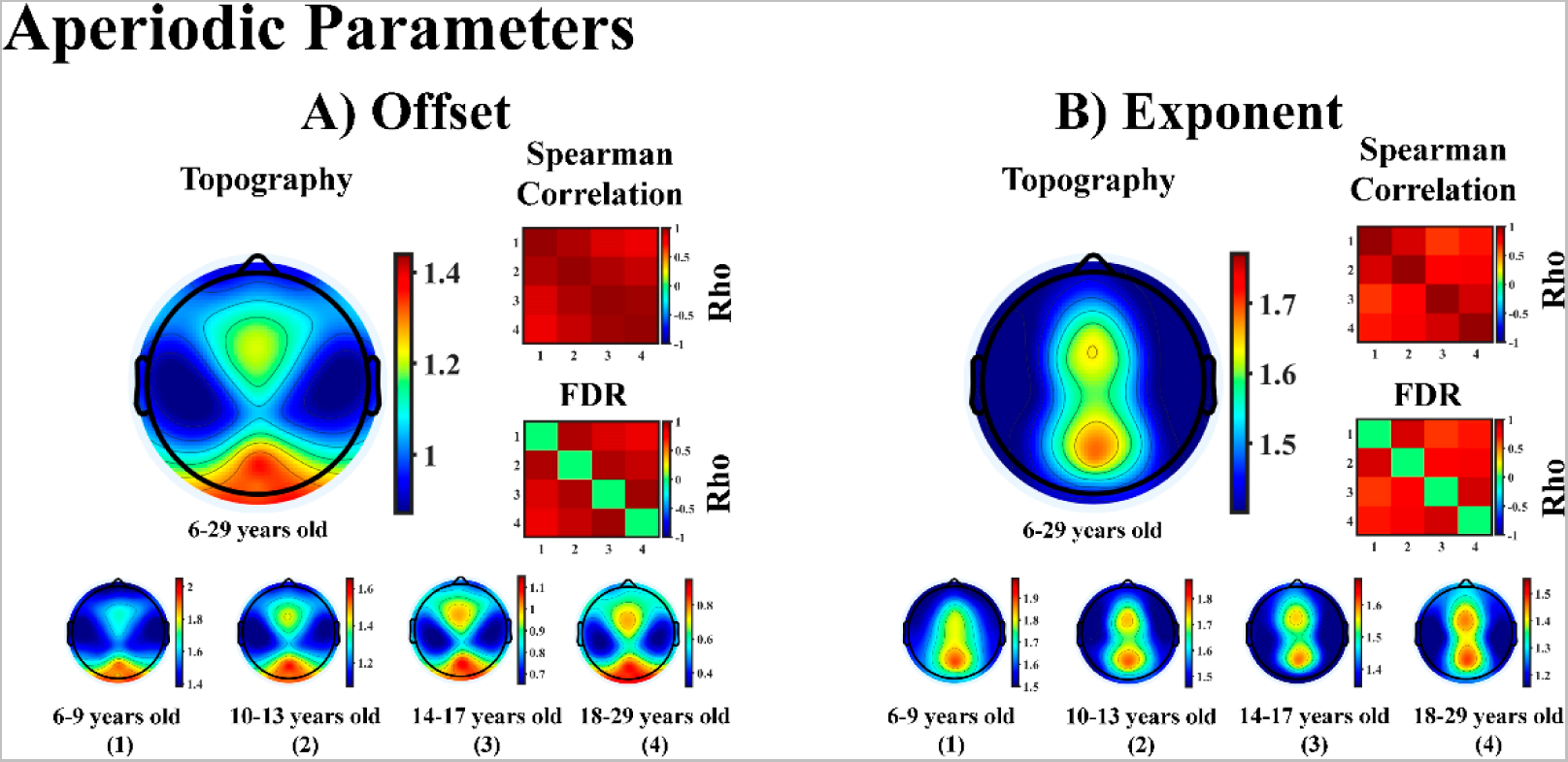
A) Topography of Offset parameter of the aperiodic PSD component in all subject and in 4 groups of age (6-9 years old, 10-13 years old, 14-17 years old and 18-29 years old). B) Topography of Exponent parameter of the aperiodic PSD component in all subject and the same 4 groups of age. Rho values indicate the Spearman correlation between the topographies of the 4 groups of offset and exponent parameters. Non-significant and diagonal row values are shown in green color (FDR corrected).

#### 2.4.3. Topographical Analysis

To observe the distribution of all PSD components on the scalp, we performed a topographical analysis. Subjects were grouped into four groups of different age ranges: 6-9 years (63 subjects), 10-13 years (59 subjects), 14-17 years (48 subjects) and 18-29 years (70 subjects). Topographies were computed in frequency windows of 3Hz in all the electrodes for the empirical PSD, PSD adjusted, AP, and P components in each of the groups. A total of 15 topographies were obtained: (1-3Hz), (4-6Hz), (7-9Hz), (10-12Hz), (13-15Hz), (16-18Hz), (19-21Hz), (22-24Hz), (25-27Hz), (28-30Hz), (31-33Hz), (34-36Hz), (37-39Hz), (40-42Hz), (43-45Hz). In addition, the offset and exponent parameters of the AP component were represented topographically.

#### 2.4.4. Spearman Correlation

Five Spearman correlation analyses were performed, each with a different purpose. When topographies are the correlated variables, Spearman correlations between the PSD values of corresponding electrodes were obtained. This method has been previously used and validated to compare the similarity of brain maps neural sources (Gómez et al., 2004, appendix). The computed correlations were: (i) The first correlation was performed to analyze the similarity between the topographies of the AP parameters (offset and exponent) between each age group; (ii) the second correlation was performed to evaluate the relationship between the empirical PSD and the PSD adjusted topographies, as a measure of the fit between the model and the empirically obtained power spectrum topographies. The topographies of the AP and P components were also computed, to determine whether or not there was a difference between the topographies of these components. If AP and P topographies were significantly correlated it would suggest similar physiological underpinnings. On the contrary, different topographies would suggest different neural sources for AP and P. This second correlation analysis was performed independently for each of the age groups.

(iii) The third correlation analysis was computed between the power of each frequency of the AP and P components. Each frequency power was correlated with all other frequency powers, for all subjects and each age group. The purpose was to observe possible similarities or differences between topographies of different frequencies. (iv) the fourth correlation analysis was performed to observe the variation of the different topographies of the PSD components (empirical PSD, PSD adjusted, AP, P) between healthy children and adults. This was motivated, first, to confirm a previous result that showed the shift towards high-frequency oscillations in the topographies from young to old (Rodríguez-Martínez et al., 2017). In fact, this is a complementary method to observe displacement to higher frequencies in a given brain area, without the need to extract a PSD peak in the power spectrum. Second, we will observe whether this shift also occurs in the different components of the PSD. Finally, (v) the objective of the fifth correlation analysis was to evaluate the relationship between all the calculated frequencies (1-45Hz) of each of the PSD components with the age (in days) of the subjects, in all electrodes, to observe age-related changes in each PSD component. The organization in frequency windows of 3Hz for the developmental trajectories of spectral power allows to compensate for the frequency shift to higher frequencies with development (Rodríguez-Martínez et al., 2017; Cellier et al., 2021). FDR was used in all correlations as a correction measure for multiple comparisons (Benjamini and Hochberg, 1995).

#### 2.4.5. Curve Estimation

To reduce the dimensionality of the data, we averaged over 15 frequency windows (see topographical analysis) the AP and P components of the spectral power. Using the Statistical Package for the Social Sciences 25 (SPSS), we performed the curve estimation analysis of a linear and quadratic model of the PSD values for the AP and P components with the age in days of the subjects. The power values of all electrodes were collapsed. As the quadratic curve estimation was superior in most cases to the linear curve, we present the results of the quadratic model. In addition, FDR was used to correct for multiple comparisons (Benjamini and Hochberg, 1995).

#### 2.4.6. Components of PSD vs. Psychological Measures

To analyze the possible relationship between the AP parameters (offset and exponent) and P component of PSD with psychological measures such as the mean reaction time (*RT*) of the Oddball task and, the subtests (Phonological Loop (*PL*), Visuospatial Sketchpad (*VSS*), Central Executive (*CE*)) of the Working Memory Test Battery for Children (*WMTBC;* Pickering and Gathercole, 2001), Spearman correlations were calculated. P-values with FDR correction (Benjamini and Hochberg, 1995) were calculated for both components (AP and P) independently.

In addition, we calculated a series of mediation analyses in JASP (2024) to determine the possible mediating role of AP and P components between age (in days) and psychological measures (RT, PL, VSS, and CE). P-values were corrected with FDR correction (Benjamini and Hochberg, 1995).

## 3. Results

Figure 1 shows different topographies for the offset and exponent parameters. The offset parameter shows a fronto-central and posterior distribution, whereas the exponent parameter shows a fronto-central and parietal distribution on the midline. The significant rho values of Spearman correlation between topographies of the different age groups ((i) correlations described in the methods section) indicate similar topographies for all age groups for the offset and exponent parameters (Figure 1).

As mentioned above, the second aim (ii) of the Spearman correlation was to analyze the degree of relationship between the empirical PSD and the model-adjusted PSD and between the AP and P components (Figures 2 and 3). The results show significant positive correlations between empirical PSD and model-adjusted PSD components in all age groups and frequency ranges (minimum Rho =0.963, p <0.001). The topographic maps of all PSD components (empirical PSD, AP and, P) as well as, for the PSD adjusted to the model are shown in Figure 2 and Figure 3. The topographies of the 15 frequency windows for each of the age groups show a similar topographical map between the empirical PSD and the adjusted PSD in all age groups. However, the topographies between the AP and P components are remarkably different in a wide range of frequencies, except for the high frequencies in which significant positive correlations are obtained, and in the middle frequencies which show negative correlations. In addition, we calculated Spearman correlations between empirical PSD and AP and P components. Empirical PSD and AP showed significant positive correlations in all age groups and frequencies (Supplementary Table 1). While correlations between empirical PSD and P show significant positive correlations around the alpha, low-beta, and gamma frequencies in all age groups (Supplementary Table 2), and significant negative correlations around high-beta and gamma, but only in adolescents and young adults.

**Figure 2.**
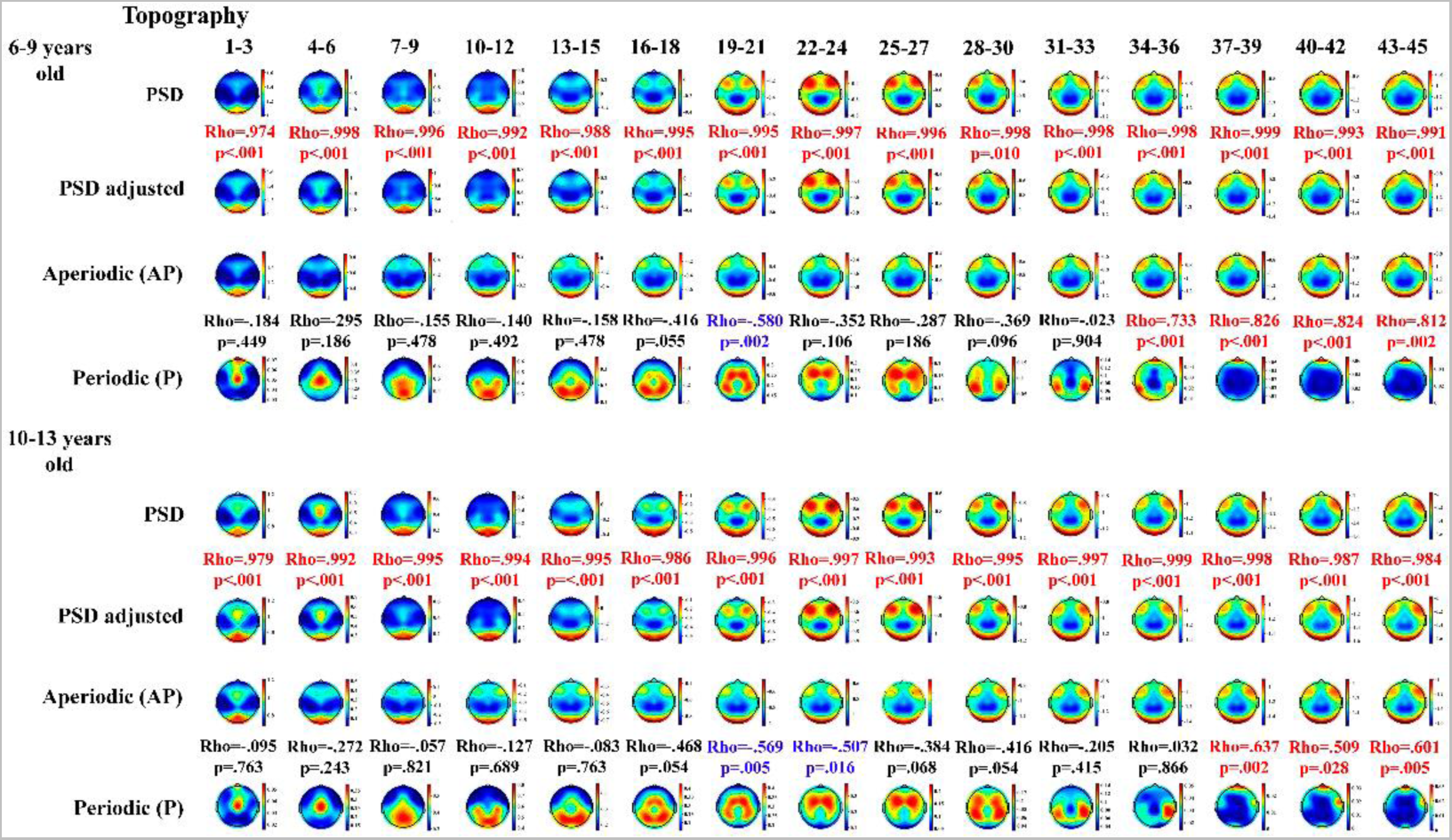
Empirical PSD, PSD adjusted, aperiodic and periodic PSD components in 6-9 and 10-13 age groups. The between maps correlation Rho values indicate the Spearman correlation (FDR corrected) between the above and below maps. The red color in the correlation indicates significant positive correlation, and blue color indicates significant negative correlation, while the black color indicates no significant correlation.

**Figure 3.**
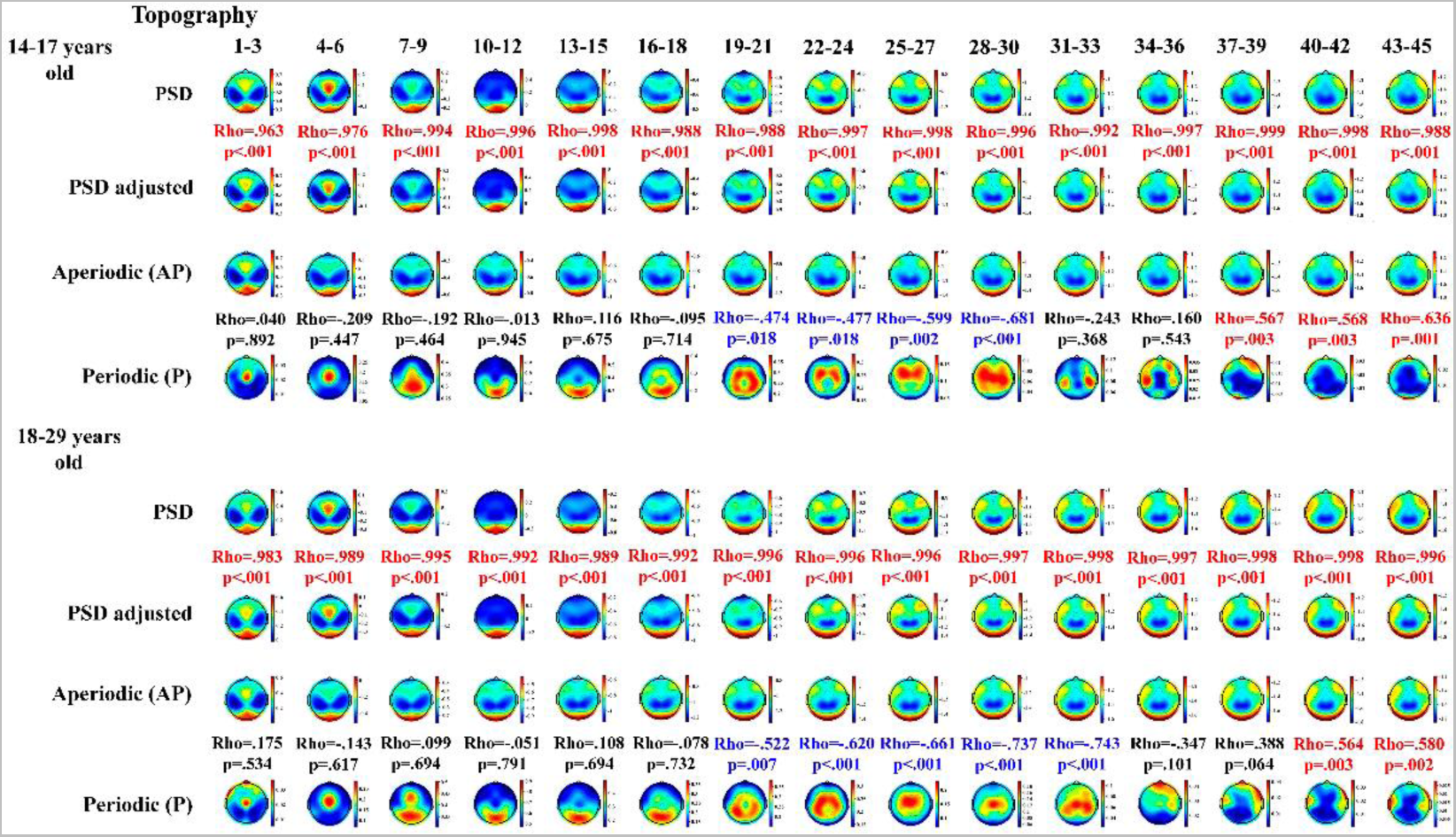
Empirical PSD, PSD adjusted, aperiodic and periodic PSD components in 14-17 and 18-29 age groups. The between maps correlation Rho values indicate the Spearman correlation (FDR corrected) between the above and below maps. The red color in the correlation indicates significant positive correlation, and blue color indicates significant negative correlation, while the black color indicates no significant correlation.

Figure 4 shows the results of the third Spearman correlation analysis. (iii) The correlation of the PSD maps across frequencies shows that AP topographies are very similar across all frequencies, also in the different age groups (inset at the right of aperiodic component). Whilst in the periodic component are theta and beta the only frequencies showing a significant similar topography (black ellipses in Figure 4), for all the age groups (inset at the right side of periodic component).

**Figure 4.**
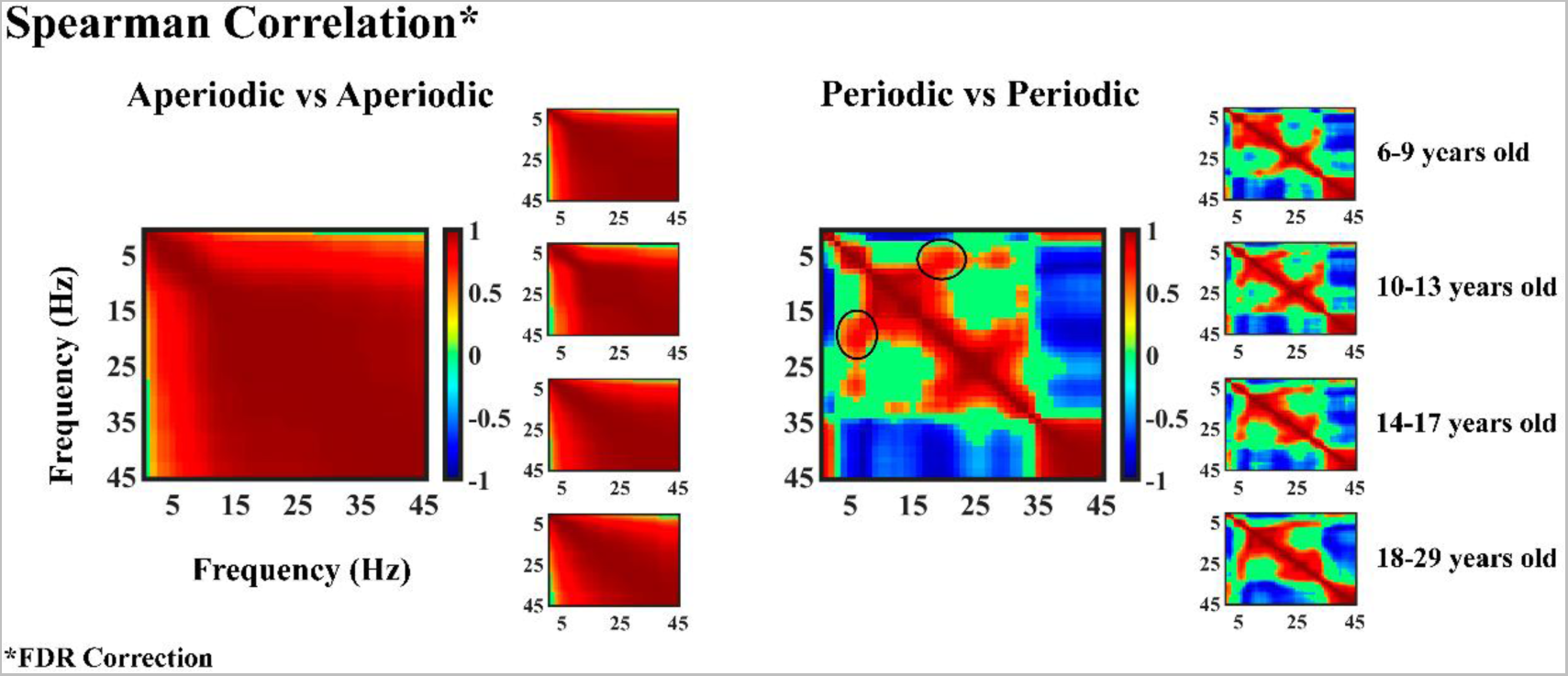
Spearman correlation of the topographies of the different frequencies of the aperiodic and periodic components of the PSD. The black ellipse shows the high co-maturation pattern of theta and beta in the periodic component topographies. Notice that correlations between topographies for the AP component was significant for almost all correlations. Green color means non-significant correlations. Statistical significance of correlations was corrected by FDR.

Figure 5 shows the results of the fourth (A) and fifth (B) proposed correlations. (iv) The fourth correlation tries to observe a possible frequency shifting in topographies depending on age (same topographies but shifted to higher frequencies as age increases). Figure 5A shows the results of the correlation analysis of the topographies of each PSD component between healthy children (6-9 years) and adults (18-29 years). There is a slight shift towards higher frequencies in the topographies of all the PSD components for adults compared to children, due to the displacement to higher frequencies of low frequencies topographies as age increases (observe the displacement of high correlations over the diagonal). The latter results suggest that children and adults present similar topographies but with a shift to lower frequencies in children, or higher frequencies in adults.

**Figure 5.**
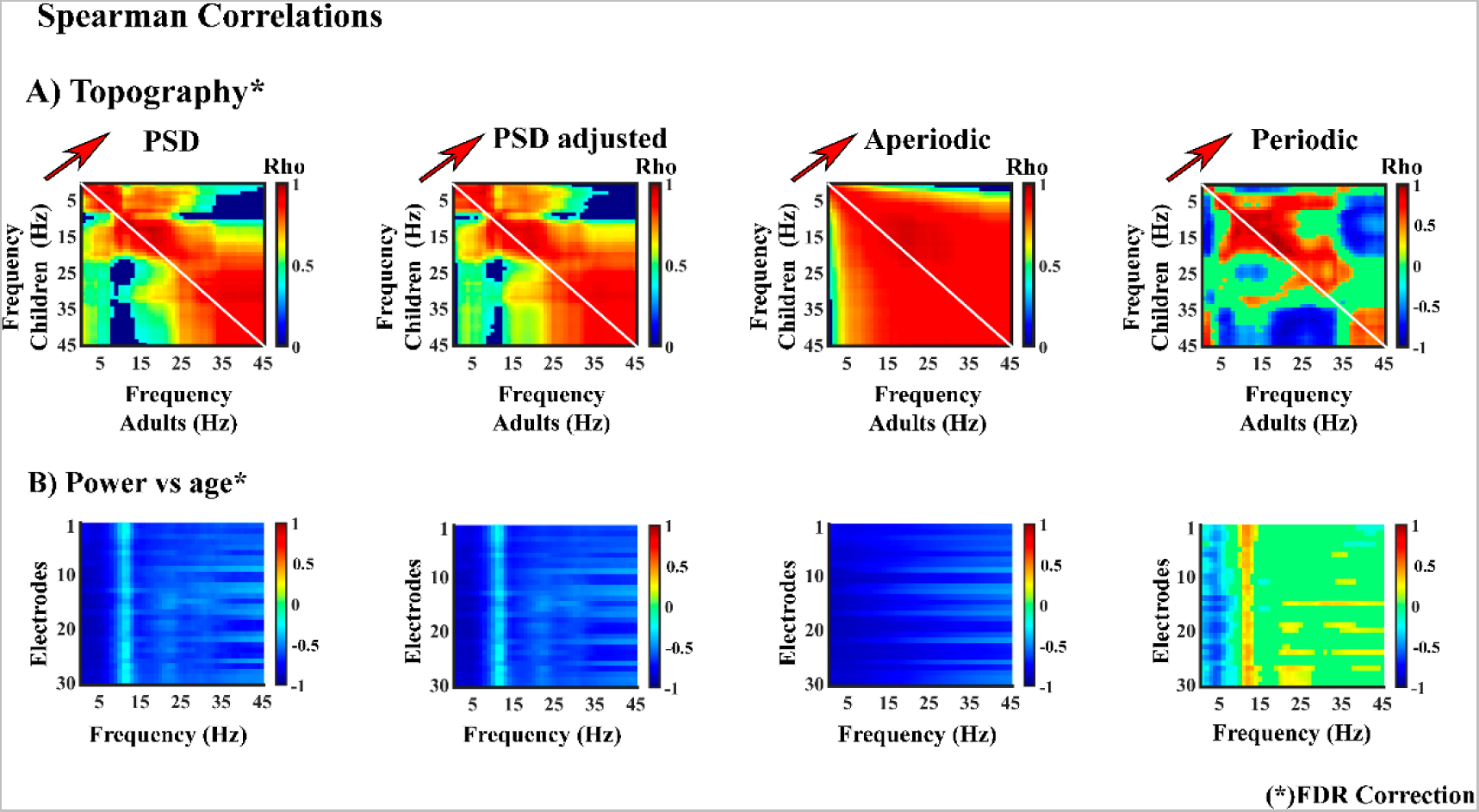
A) Spearman correlation of topographies of children vs. adults for all PSD components (in the range of 1-45Hz). The Y-axis corresponds to the children group (6-9 years old) and on the X-axis to the adult group (18-29 years old). The diagonal line indicates the location of correlation between topographies of both groups for the same frequency. The red arrow indicates the trend of change to higher frequencies for adults (similar topography but with a shift to higher frequencies of adults with respect to children). 0 values in rho indicate no significant correlation. B) Spearman correlation between the PSD of the different components with the age in days of subjects. On the X-axis, frequency range of 1-45Hz and on the Y-axis all of the electrodes analyzed (1-30). Green color in the panels indicate non-significant correlations after FDR correction.

(v) To observe the general landscape of the relationship between the PSD components with age, a Spearman correlation of PSD values (empirical PSD, PSD adjusted, AP, and P) for all the electrodes and frequencies were computed. The results of the correlation between all the calculated frequencies (1-45Hz) of each of the PSD components with age (in days) show a predominance of negative correlations in all the components except for the P component, where there are positive correlations in frequencies higher than 10Hz (see Figure 5B).

The results of the quadratic curvilinear estimation analysis between the AP parameters (offset and exponent) vs. age show a general decrease with increasing age (Figure 6A). Both offset and exponent parameters were averaged across all electrodes for each subject to reduce data dimensionality. Additionally, a linear relationship between offset and exponent AP parameters (R^2^=.790, p<.001) was obtained (Figure 6B).

**Figure 6.**
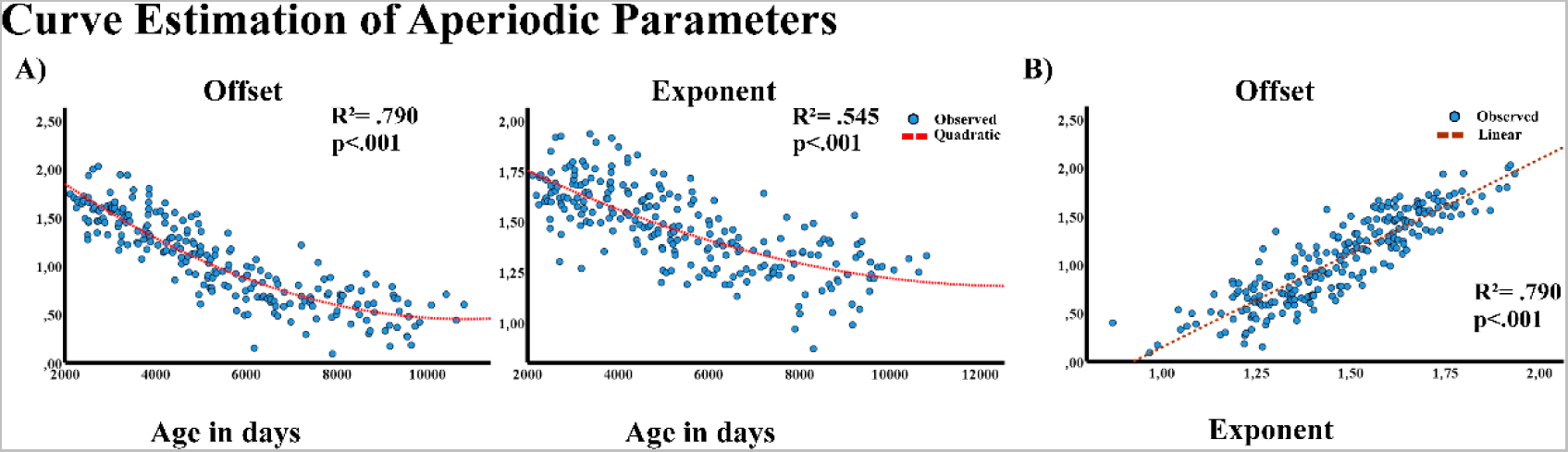
A) Quadratic regression of the parameters of the aperiodic PSD component (offset and exponent) and the age of the subjects (in days). B) Linear regression of the exponent and offset parameters. Please notice that the offset and exponent parameters have been collapsed for all the electrodes.

To reduce dimensionality and refine the type of model relating AP PSD with age, the quadratic regression of AP PSD collapsing all the electrodes, and organizing AP PSD in 15-frequency windows was computed (Figure 7). Results show a significant fitting to the quadratic curvilinear model, indicating a quadratic decrease in AP PSD as age increases.

**Figure 7.**
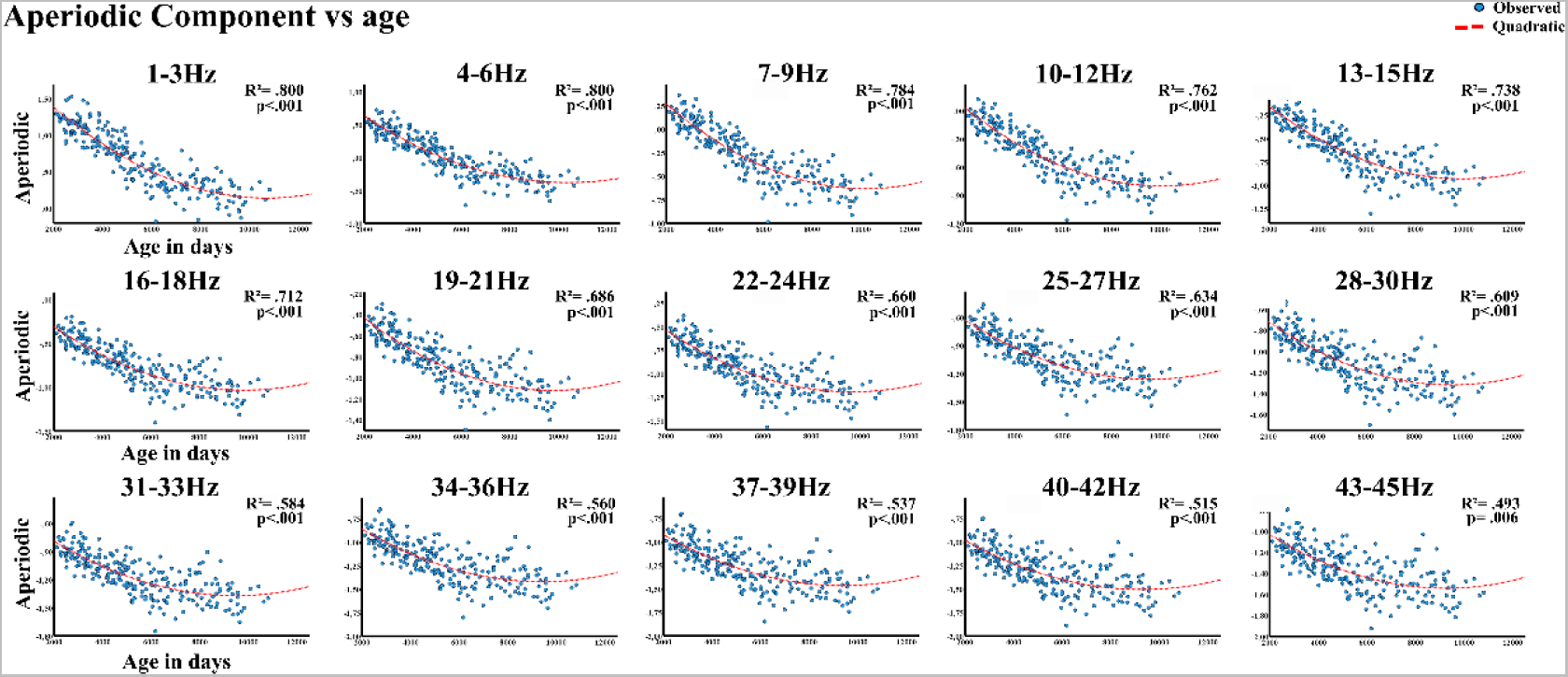
Quadratic regression model of the aperiodic PSD component vs. age (expressed in days). Aperiodic PSD values were collapsed over frequency windows of 3Hz (1-45Hz) and, across all electrodes.

Figure 8A shows the P PSD in the different age groups. As with the AP PSD component, the data of the P PSD component vs age was fitted with a quadratic curvilinear model and collapsed across electrodes. However, only the 1-3Hz, 4-6Hz, 10-12Hz, and 13-15Hz frequency bands were significant (FDR corrected) (Figure 8B).

**Figure 8.**
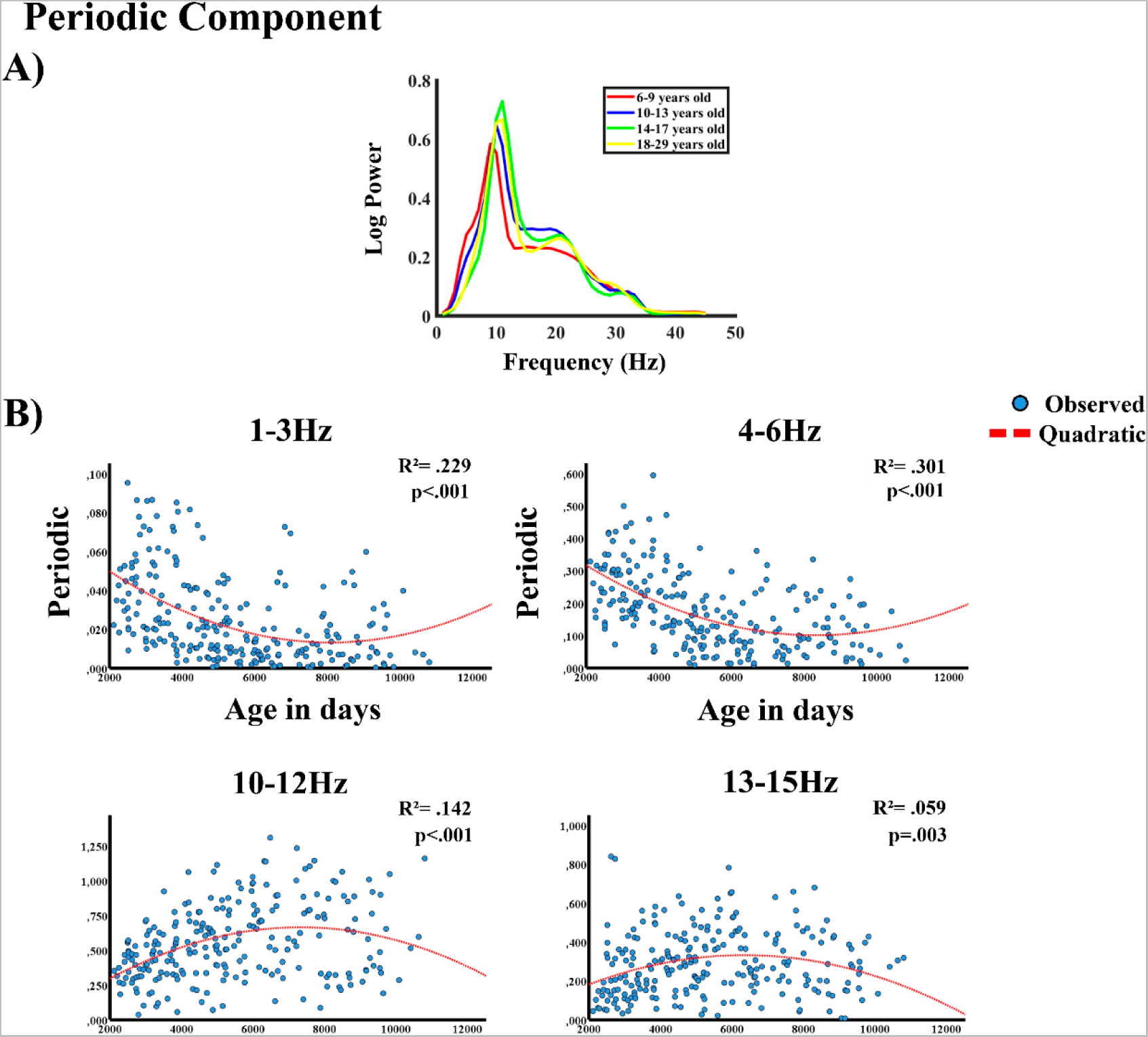
A) Periodic component of the PSD in all age ranges. B) Quadratic regression model of the periodic PSD component vs. age (expressed in days). Periodic PSD values were collapsed over frequency windows of 3Hz (1-45Hz) and, across all electrodes. Only significant regressions are displayed.

Finally, the Spearman correlation of AP parameters (offset and exponent) and P PSD vs. psychological measures (see methods and Table 2), were computed. P and AP PSD values were collapsed across electrodes and in frequency ranges of 3Hz. The results show a significant Spearman correlation for all psychological measures with the AP parameters (offset and exponent). Age-related P PSD frequency ranges (1-3Hz, 4-6Hz, 10-12Hz, 13-15Hz) also showed significant correlations with the psychological measures, except for the frequency range 13-15Hz vs. RTs and the central executive component of Working Memory. The results of the mediation analysis (Table 3), show that age effects on the RT, VSS, and CE were mediated by the offset parameter. Furthermore, the RT was also mediated by the P PSD of the theta rhythm (4-6Hz).

**Table 2.**
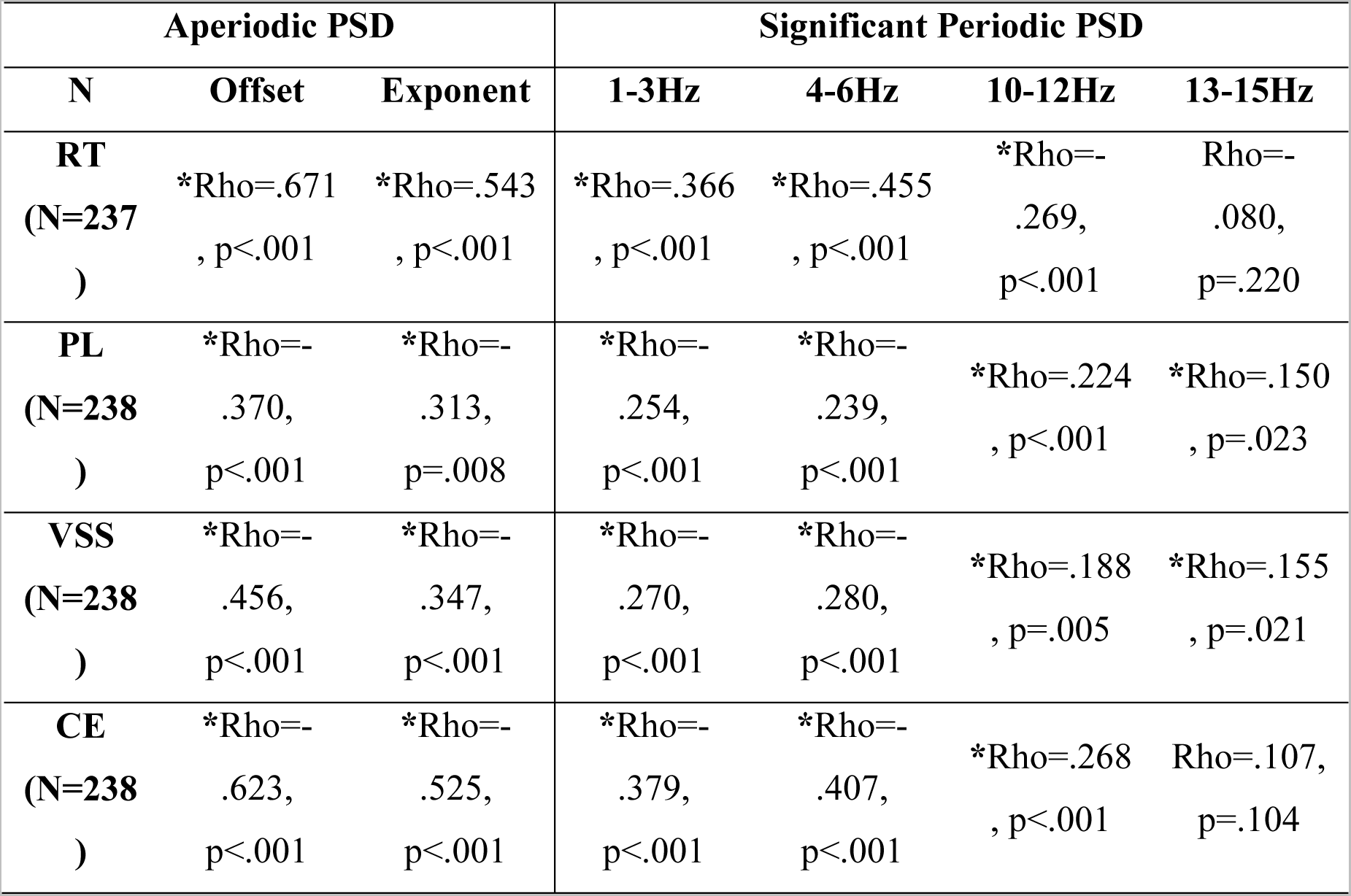
Spearman Correlation between aperiodic parameters (offset and exponent) and age-related periodic PSD vs. mean reaction time (RT) of the Oddball task, the subtests of the Working Memory Test Battery for Children (WMTBC). Phonological Loop (PL), Visuospatial Sketchpad (VSS) and Central Executive (CE). Asterisk (*) indicates a significant relationship (FDR corrected).

**Table 3.**
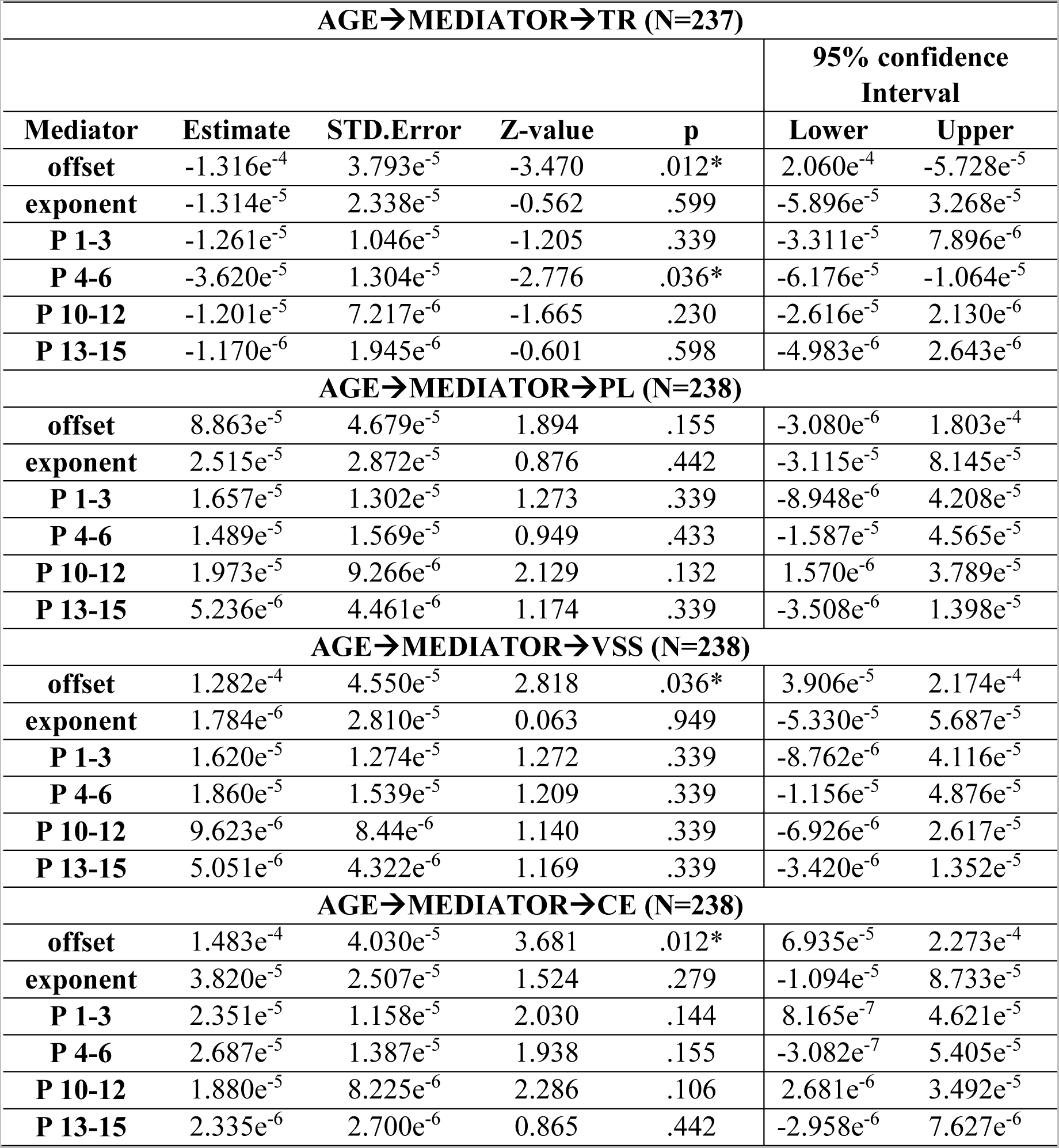
Indirect effects computed by Mediation Analysis of the aperiodic parameters (offset and exponent) on the age vs. psychological measures. The indirect effects of the age-related periodic PSD values on the age vs. psychological measures are also displayed. P-values with FDR correction. Significant p-values with asterisk (*).

## 4. Discussion

In this study, we analyzed the AP and P components of spectral power during resting state in a large group (N=240) and a broad age range (6-29 years old) of typical healthy subjects. Previous research suggests that the AP and P components of the EEG signal are dynamically and physiologically distinct (Donoghue et al., 2020a; He, 2014; Voytek and Knight, 2015). There was a clear topographical difference between the AP and P components suggesting different neural sources. The differential developmental trajectory of AP and P PSD also suggests the neurophysiological independence of AP and P. On the other hand, the displacement to higher frequencies of adult topographies obtained in the present report (AP and P), suggests that the well-described displacement to higher frequencies in the alpha peak (Cellier et al., 2021; Donoghue et al., 2020a; Hill et al., 2021; McSweeney et al., 2023) extends to other frequencies above and below the dominant frequency. The co-maturation of P and AP components with WM-related measures extends the previously described developmental cognitive value of AP and P components, being the offset AP component the most robust mediator between age and overt behavior.

We analyzed the topographic distribution of the AP component parameters (offset and exponent) and different topographic distributions were observed (Hill et al., 2022; Jacob et al., 2021). The offset and exponent topographies were maintained throughout maturation. The offset parameter shows a fronto-central and posterior topographic distribution, with higher posterior values. The exponent is mainly distributed over fronto-central and parietal areas on the midline of the scalp and presents higher posterior values. These findings confirm recent studies in EEG (Hill et al., 2022; Jacob et al., 2021) and MEG (Donoghue et al., 2020a).

The topographies of P and AP PSD components were quite different, only high frequencies presented a similar topography between P and AP, and given the constancy across frequencies of the AP component, the gamma range of the PSD would be dominated by the AP component. Regarding the P component, although the current study examines it without defining *a priori* frequency bands, it shows regionally delineated oscillations partially coincident with regions described in the literature for canonical frequency ranges, such as delta (0.5-3Hz), theta (4-7Hz), alpha (7-14Hz) and beta (15-30Hz) (Donoghue et al., 2020a; see Fig.1 in Gómez et al., 2006; Rodríguez-Martínez et al., 2012, 2017, 2020). Similar patterns have been observed in magnetoencephalography (MEG) (Donohue et al., 2020a; Jacob et al., 2021) and electroencephalography (Hill et al., 2022) studies, showing power concentration for delta at frontal areas, theta at the central midline, alpha in posterior areas, beta centered on sensorimotor areas and gamma bi-temporally (Jacob et al., 2021; Niso et al., 2016). However, a more detailed topographical correlational analysis suggests that canonical topographies are more dominated by the AP PSD component than by the P PSD component, except for alpha and low-beta in which the topography correlational analysis suggests an important contribution of P PSD. Taken together, all the previously described results suggest that canonical EEG bands during resting state share similar topographies with the P component for alpha, low-beta, and gamma. Nevertheless, the P PSD component is embedded in the canonical topography for delta, theta, and high-beta. The latter results suggest that the myriad of previous results obtained with the canonical approach are still valid, although they can be refined by the parameterization approach by discounting the AP influence on the PSD.

Some studies of EEG oscillations suggest regional changes in theta oscillations with age, considering posterior theta as a precursor of posterior alpha, (Barriga-Paulino et al., 2011; Cellier et al., 2021), but also the presence of an independent emergent fronto-central topography across development (Cellier et al., 2021) has been described. Our data support these results, particularly for the alpha range, although the presence of a central theta oscillation in the younger group suggests a relative early emergence of central theta. Additionally, the increase in frequency of oscillations as detected in the transition from posterior theta to alpha, can be extended to the whole spectral frequency range, as revealed by the change in topographies towards higher frequencies with age for all components (empirical PSD, PSD adjusted, AP and P), as it was previously shown in studies of the canonical PSD (Rodríguez-Martínez et al., 2012, 2017). In fact, the displacement to higher frequencies of the oscillations in both AP and P as revealed by the correlation of topographies of different ages, suggests that the increases in the frequency of oscillations in the cortex are a phenomenon that extends the dominant frequency (Cellier et al., 2021; Donoghue et al., 2020a; Hill et al., 2021; McSweeney et al., 2023; Rodríguez-Martínez et al., 2017). The studies that have shown changes with age in aperiodically adjusted frequencies mainly analyze the maximum frequency of dominant oscillations. Thus, most of these studies only show increases with age of the maximum frequency of the dominant alpha oscillation (8-12Hz) (Cellier et al., 2021; He et al., 2019; Hill et al., 2022). Recently a study also shows a decrease with age of the peak frequency of periodic beta in the age range of 4 to 10 years, with a stabilization of peak frequency around 8 years (McSweeney et al., 2023). The present report suggests that for the beta range, there is a displacement of oscillatory frequency in the transition to adulthood, at least when a topographical approach is implemented.

For age-related changes in power in the AP parameters and power, decreases with age have been previously observed (Cellier et al., 2021; Dave et al., 2018; Donoghue et al., 2020a; Hill et al., 2022; McSweeney et al., 2021, 2023) and these changes have been linked to neuroanatomical (He et al., 2019; Jacob et al., 2021) and neurochemical (Buzsáki et al., 2012; Cohen Kadosh et al., 2015) changes. While our results align with these findings: decreases of AP PSD with age at all electrodes and frequencies, we emphasize the presence of a quadratic explanatory model with maturation as it has been recently proposed (McSweeney et al., 2023). Canonical PSD EEG studies differ in their trend during maturation depending on the power analysis performed (absolute PSD or relative PSD). As several authors have argued, absolute PSD (Gasser et al., 1988; Miskovic et al., 2015; Rodríguez-Martínez et al., 2012, 2017; Segalowitz et al., 2010) decreases with age, while the variability increases (Angulo-Ruiz et al., 2021, 2022), which has been attributed to the natural maturation processes of the nervous system, such as neural pruning (Whitford et al., 2007). In this regard, our results emphasize that the decrease in absolute PSD would be mostly due to a decrease in the AP component with age, as reported in other studies (Cellier et al., 2021; Dave et al., 2018; Donoghue et al., 2020a, 2020b; He et al., 2019; Hill et al., 2022; McSweeney et al., 2021, 2023; Schaworonkow and Voytek, 2021) since it appears at all frequencies. Changes with age would not only be due to neural pruning (Whitford et al., 2007) but also to structural decreases in cortical thickness/volume (Giedd et al., 1999; Giedd, 2004; Gogtay et al., 2004; Paus, 2005), changes in the recruited neuronal population (Manning et al., 2009; Miller et al., 2014) and changes in the balance between excitatory and inhibitory neurotransmitters (Cohen Kadosh et al., 2015; Donoghue et al., 2020a; Gao et al., 2017; Waschke et al., 2021). All critical for optimal cortical development and function (Donoghue et al., 2020a). The correlation of topographies of different frequencies showed that in the P component theta and beta topographies were the only positive correlations, while in the AP component topographies were significantly similar for almost all analyzed frequencies (1-45Hz). The latter results suggest that AP component topography would be dependent on anatomical constraints, on how the electrical dipolar activity generating the AP activity projects on the scalp.

EEG power can also be characterized as relative power. Most studies of relative PSD indicate a greater contribution to higher frequencies to the total spectral energy as age increases (Segalowitz et al., 2010; Rodríguez-Martínez et al., 2017) and, would suggest the shift in connectivity trends, with an increase in local connectivity (fast frequencies) over long-range connectivity (slow frequencies) (Lea-Carnall et al., 2016), as well as an increase in the velocity of action potentials due to myelination and/or increased axon diameter (Barriga-Paulino et al., 2011; Segalowitz et al., 2010; Sowell et al., 2002). The redistribution of relative PSD would be partially a consequence of the AP exponent value decrease with age, which would flatten the EEG power spectrum. The latter would suggest a change in the balance of excitatory (E) and inhibitory (I) synaptic currents (Buzsáki et al., 2012; Gao et al., 2017; Waschke et al., 2021). In this sense, a lower exponent value would signify a higher excitatory than inhibitory current at the cortical level (Donoghue et al., 2020a; Gao et al., 2017). This increase in excitatory activity in the cortex could also explain neuroplasticity and the acquisition of new cognitive skills as age increases (Cohen Kadosh et al., 2015; Dave et al., 2018; Donoghue et al., 2020a).

The present report analyzes the P component by aperiodically adjusting the obtained oscillations and relates them to age. Age-related changes in the P component, with a more restricted frequency range than in the AP, were observed. Therefore, changes with age in the power of P oscillations are supported by the present results (Cellier et al., 2021; Donoghue et al., 2020a; He et al., 2014; Hill et al., 2022; McSweeney et al., 2023). EEG P power changes are mainly observed in the 1-15Hz range with decreases in the lower range (1-6Hz) and increases in the higher range (10-15Hz). This is in agreement with recent studies reporting decreases in the theta band power with age (McSweeney et al., 2023) and increases in the alpha band (Cellier et al., 2021; Donoghue et al., 2020a; Hill et al., 2022; McSweeney et al., 2023), bands that coincide with the ranges considered in this study. We also observed power increases with age in the beta P range (20-27Hz), (He et al., 2019; McSweeney et al., 2023), but restricted to the posterior electrodes. We confirm that the power of oscillations in theta, alpha, and beta ranges changes with age. Furthermore, we also suggest that this change extends to the delta frequency band (1-3Hz), where decreases with age are observed. In summary, we show that characteristics of P oscillations in different frequency ranges are associated with age, with specific patterns of change occurring over time.

There is a growing body of research showing that some cognitive processes are related to the AP component, including attention (Mamiya et al., 2021), task performance (Donoghue et al., 2020a; Ostlund et al., 2021), processing speed (Ostlund et al., 2021; Ouyang et al., 2020), sensory processing (Tran et al., 2020), language (Dave et al., 2018), or working memory (Donoghue et al., 2020a; Gao et al., 2017). This relationship is explained by a decrease in the exponent AP parameter and changes in excitatory (E) and inhibitory (I) neuronal currents, produced as a result of an increase in the ratio of glutamate (E) to GABA (I) (Cohen Kadosh et al., 2015; Donoghue et al., 2020a; Gao et al., 2017; Waschke et al., 2021). Some research has shown how high levels of glutamate in specific brain regions (e.g., inferior frontal gyrus) positively correlate with enhanced cognitive processing, such as proficiency in face processing (Cohen Kadosh et al., 2015), or attentional control (Mamiya et al., 2021). Our results are in line with this approach and suggest covariations between the AP component and psychological variables of processing speed (RT) and memory (PL, VSS, CE). However, the mediational effect for the relationship between age and behavior is exerted by the offset parameter, in contrast to studies pointing to a higher effect of the AP exponent on behavior (Ouyang et al., 2020; Podvalny et al., 2015; Voytek et al., 2015b; Waschke et al., 2021). The latter result suggests that different behavioral and cognitive processes can be influenced by different AP parameters across development. Therefore, more neurophysiological studies with psychological correlates are needed to argue that changes in the AP activity of the EEG signal are the result of changes in the structure and organization of the brain during development.

In summary, our results emphasize the significance of PSD parameterization for a more effective interpretation of attributions related to neural oscillations in various physiological and cognitive processes (Donoghue et al., 2020a; Ostlund et al., 2022). This approach avoids confusion of the genuine oscillatory (P) activity with the AP component, which is arrhythmic in nature (Donoghue et al., 2020a; He, 2014). However, this study has some limitations, and the results should be interpreted cautiously. The main limitation is that since our study focused on the analysis of AP PSD activity as a function of its parameters (exponent and offset), differential analysis of P activity for its parameters (center frequency, power, and bandwidth) was not considered, instead focusing on power and topographic correlations. Furthermore, since our main interest lies in changes during maturation, we excluded other relevant variables that have been shown to influence AP activity, such as gender (McSweeney et al., 2021), after showing no effects on behavior and AP parameters. It should also be noted that our study uses a cross-sectional sample, and for characterizing individual differences, it would be useful to analyze these components longitudinally.

## 5. Conclusions

The parameterization of spectral power into distinct components enhances the interpretation, and neurophysiological and functional accuracy of brain processes. Our results support the existence of different PSD components that change dynamically with maturation and clarify the relationship with physiological and psychological processes described in the literature.

## Statements and declarations

### Competing Interest

None of the authors have potential conflicts of interest to be disclosed.

## Supporting information

Supplementary Figura and Tables

## Acknowledgements

We are grateful to the children, adolescents, and young adults who participated in the present study.

## Financial Support

This study was supported by grants from the Agencia Estatal de Investigación [PID2019– 105618RB-I00 and PID2022-139151OB-I00]; and the Agencia de Innovación y Desarrollo de la Junta de Andalucía [P20_00537].

## Notes

### Competing Interest Statement

The authors have declared no competing interest.

