## Supplementary Figura and Tables for "Unveiling the hidden electroencephalographical rhythms during development: aperiodic and periodic activity in healthy subjects"

**SUPLEMENTARY FIGURE**

Vanesa Muñoz[1]

Carlos M. Gómez[1],

Corresponding author:

Carlos M. Gómez[1]

[1] Human psychobiology laboratory, Experimental Psychology Department, University of Seville, C/ Camilo José Cela S/N 41018 Seville, Spain.

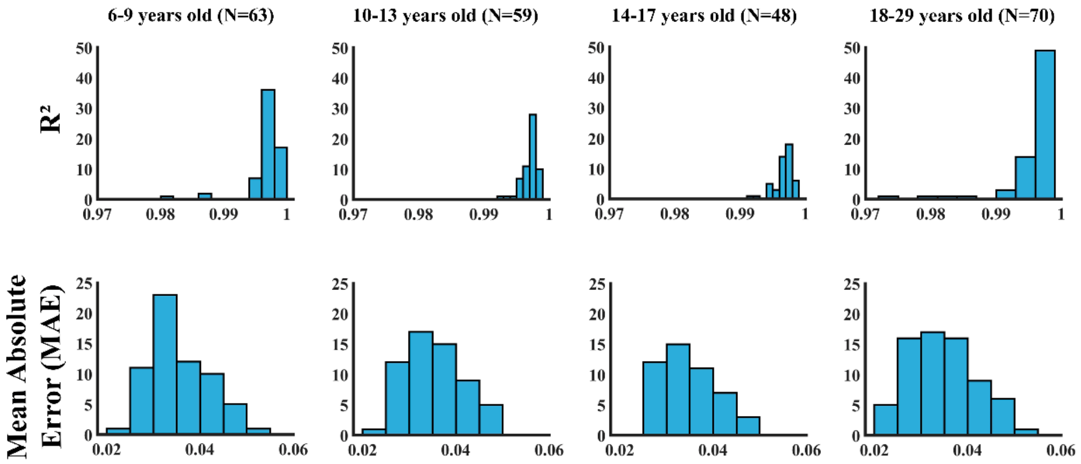


**Supplementary Figure 1.** Histograms of explained variance ($R^2$) and mean absolute error (MAE) metrics to assess the goodness-of-fit measures between model and original power spectrum.

**SUPLEMENTARY TABLES**

### Unveiling the hidden electroencephalographical rhythms during development: aperiodic and periodic activity in healthy subjects

Brenda Y. Angulo-Ruiz[1],

Elena I. Rodríguez-Martínez[1],

Vanesa Muñoz[1]

Carlos M. Gómez[1],

Corresponding author:

Carlos M. Gómez[1]

[1] Human psychobiology laboratory, Experimental Psychology Department, University of Seville, C/ Camilo José Cela S/N 41018 Seville, Spain.

**Supplementary Table 1**

Spearman Correlation between empirical PSD and Aperiodic Component topographies for all age groups. P-values with FDR correction. Please notice that all correlations were significant.

| | 6-9 years old | | 10-13 years old | | 14-17 years old | | 18-29 years old | |
|---|---|---|---|---|---|---|---|---|
| **Frequencies** | **Rho** | **p** | **Rho** | **p** | **Rho** | **p** | **Rho** | **P** |
| **1-3** | .995 | <.001 | .960 | <.001 | .962 | <.001 | .980 | <.001 |
| **4-6** | .812 | <.001 | .857 | <.001 | .867 | <.001 | .918 | <.001 |
| **7-9** | .539 | .002 | .717 | <.001 | .752 | <.001 | .810 | <.001 |
| **10-12** | .484 | .007 | .330 | .075 | .496 | .005 | .511 | .003 |
| **13-15** | .700 | <.001 | .572 | .001 | .648 | <.001 | .660 | <.001 |
| **16-18** | .774 | <.001 | .624 | <.001 | .720 | <.001 | .735 | <.001 |
| **19-21** | .873 | <.001 | .728 | <.001 | .725 | <.001 | .717 | <.001 |
| **22-24** | .884 | <.001 | .797 | .001 | .826 | <.001 | .834 | <.001 |
| **25-27** | .947 | <.001 | .911 | <.001 | .948 | <.001 | .929 | <.001 |
| **28-30** | .973 | <.001 | .969 | <.001 | .953 | <.001 | .941 | <.001 |
| **31-33** | .965 | <.001 | .972 | <.001 | .965 | <.001 | .960 | <.001 |
| **34-36** | .996 | <.001 | .993 | <.001 | .997 | <.001 | .989 | <.001 |
| **37-39** | .998 | <.001 | .989 | <.001 | .995 | <.001 | .993 | <.001 |
| **40-42** | .992 | <.001 | .980 | <.001 | .994 | <.001 | .995 | <.001 |
| **43-45** | .990 | <.001 | .977 | <.001 | .983 | <.001 | .993 | <.001 |

**Supplementary Table 2**

Spearman Correlations between empirical PSD and Periodic Component topographies for all age groups. P-values with FDR correction. Significant correlations (negative or positive) with asterisk (*).

| | 6-9 years old | | 10-13 years old | | 14-17 years old | | 18-29 years old | |
|---|---|---|---|---|---|---|---|---|
| **Frequencies** | **Rho** | **p** | **Rho** | **p** | **Rho** | **p** | **Rho** | **P** |
| **1-3** | -0.34 | .856 | -.015 | .954 | .152 | .489 | .280 | .168 |
| **4-6** | .259 | .313 | .197 | .495 | .182 | .458 | .162 | .408 |
| **7-9** | .657 | <.001* | .600 | .001* | .457 | .021* | .634 | <.001* |
| **10-12** | .721 | <.001* | .813 | .001* | .791 | .006* | .746 | <.001* |
| **13-15** | .527 | .006* | .699 | <.001* | .777 | .011* | .726 | <.001* |
| **16-18** | .228 | .357 | .481 | .046* | .564 | .004* | .528 | .005* |
| **19-21** | -.158 | .505 | .086 | .863 | .221 | .362 | .166 | .408 |
| **22-24** | .053 | .838 | .011 | .954 | .036 | .849 | -.157 | .408 |
| **25-27** | -.056 | .838 | -.074 | .863 | -.386 | .059 | -.395 | .046* |
| **28-30** | -.222 | .357 | -.268 | .285 | -.478 | .016* | -.539 | .005* |
| **31-33** | .173 | .492 | -.061 | .863 | -.095 | .663 | -.597 | .002* |
| **34-36** | .748 | <.001* | .084 | .863 | .168 | .47 | -.284 | .168 |
| **37-39** | .830 | <.001* | .689 | <.001* | .596 | .003* | -.417 | .036* |
| **40-42** | .862 | <.001* | .598 | .004* | .637 | .002* | .583 | .002* |
| **43-45** | .853 | <.001* | .686 | <.001* | .639 | <.001* | .612 | .001* |
